# Transcriptome Analysis of *Neisseria gonorrhoeae* During Natural Infection Reveals Differential Expression of Antibiotic Resistance Determinants Between Men and Women

**DOI:** 10.1101/319970

**Authors:** Kathleen Nudel, Ryan McClure, Matthew Moreau, Emma Briars, A. Jeanine Abrams, Brian Tjaden, Xiao-Hong Su, David Trees, Peter A. Rice, Paola Massari, Caroline A. Genco

## Abstract

*Neisseria gonorrhoeae* is a bacterial pathogen responsible for the sexually transmitted infection, gonorrhea. Emergence of antimicrobial resistance (AMR) of *N. gonorrhoeae* worldwide has resulted in limited therapeutic choices for this infection. Men who seek treatment often have symptomatic urethritis; in contrast, gonococcal cervicitis in women is usually minimally symptomatic, but may progress to pelvic inflammatory disease. Previously, we reported the first analysis of gonococcal transcriptome expression determined in secretions from women with cervical infection. Here, we defined gonococcal global transcriptional responses in urethral specimens from men with symptomatic urethritis and compared these with transcriptional responses in specimens obtained from women with cervical infections, and of *in vitro-grown N. gonorrhoeae* isolates. This is the first comprehensive comparison of gonococcal gene expression in infected men and women. RNA sequencing analysis revealed that 9.4% of gonococcal genes showed increased expression exclusively in men and included genes involved in host immune cell interactions and 4.3% showed increased expression exclusively in women and included phage-associated genes. Infected men and women displayed comparable antibiotic resistant-genotypes and *in vitro* –phenotypes, but a 4-fold higher expression of the Mtr efflux pump-related genes was observed in men. These results suggest that expression of AMR genes is programmed genotypically, and also driven by sex-specific environments. Collectively, our results indicate that distinct *N. gonorrhoeae* gene expression signatures are detected during genital infection in men and women. We propose that therapeutic strategies could target sex specific differences in expression of antibiotic resistance genes.

## Importance

Recent emergence of antimicrobial resistance of *Neisseria gonorrhoeae* worldwide, has resulted in limited therapeutic choices for treating infections caused by this organism. We performed global transcriptomic analysis of *N. gonorrhoeae* in subjects with gonorrhea who attended the Nanjing (China) STI clinic where antimicrobial resistance of *N. gonorrhoeae* is high and increasing. We found that *N. gonorrhoeae* transcriptional responses to infection differed in genital specimens taken from men and women, particularly antibiotic-resistance gene expression, which was increased in men. These sex-specific findings may provide a new approach to guide therapeutic interventions and preventive measures that are also sex-specific while also providing additional insight to address antimicrobial resistance of *N. gonorrhoeae*.

## Introduction

*Neisseria gonorrhoeae* is responsible for the sexually transmitted infection (STI), gonorrhea. Rates of gonococcal infection in the United States have increased 46% since 2011, and approximately 450,000 cases of gonorrhea were reported to the U.S. Centers for Disease Control and Prevention in 2015. *N. gonorrhoeae* is the second most common reportable bacterial STI in the U.S. (1). Infection with *N. gonorrhoeae* also contributes significantly to global STI morbidity and is responsible for 78 million cases each year (2). For the past eight decades, gonorrhea has been treated successfully with antibiotics but over the last several years antibiotic resistance has begun to emerge worldwide. Resistant strains include isolates from both Eastern and Western Europe (3, 4), Asia (5–7) and recently the United States (8–10). This rise in antibiotic resistant strains may be linked, in part, to an increase in availability of over-the-counter antibiotics, particularly in many parts of Asia. New strategies for combatting this disease are necessary, as evidenced by increased efforts to develop gonococcal vaccines (11, 12) and new drug regimens (7, 13, 14).

In humans, the only natural host for *N. gonorrhoeae*, symptomatic responses to infection are unique to men and women. While most infected men remain asymptomatic (15), those who develop symptoms often show robust inflammation characterized by purulent urethral discharge accompanied by large numbers of polymorphonuclear leukocytes (PMNs). In women, gonococcal infections are asymptomatic 50–80% of the time (16) or are accompanied by non-specific symptoms such as vaginal discharge (17). The presence of cervical mucus or microscopic PMN counts ≥10 seen microscopically on an oil immersion field does not correlate well with gonococcal infection in the absence of co-infecting agents (18). Manifestations of gonococcal infection may be driven by environmental factors specifically associated with the genital tracts of men and women: these may include biofilm formation (19, 20); the influence of the microbiome, particularly in women (21–25) and both direct and indirect molecular interactions between gonococci and specific host cells from men or women (26–30).

Gonococcal pathogenesis and host immune responses have been studied principally *in vitro* and in animal models (31), both of which may not faithfully replicate differences in infection between men and women. Historically, the only human model of gonococcal infection is based on genital experimental challenge of male volunteers, which has provided valuable information on gonococcal virulence factors, but is limited by the duration of the experimental infection and is not applicable to women (32, 33). A better understanding of gonococcal pathogenesis during natural mucosal infection, which is less well studied, is critical to implement biologic strategies to treat and prevent infection. Previous work from our group using a gene specific microarray and qRT-PCR analysis demonstrated that a subset of gonococcal iron-regulated genes were expressed during human mucosal infection in men and women (34, 35). Recently, we also examined the complete *N. gonorrhoeae* transcriptome using cervico-vaginal lavage specimens of infected women attending an STI clinic in China where antibiotic resistant *N. gonorrhoeae* is prevalent (36, 37). We reported that a large portion of the gonococcal genome is expressed and regulated during infection (38). In the present study, we examined gonococcal transcriptomes from men attending the same clinic and compared bacterial gene expression and regulation in infected men with expression and regulation in infected women. The resulting data revealed sex-specific adaptation of genes involved in host-pathogen interactions, phage and antibiotic resistance.

## Results

### Description of Subjects

The male cohort comprised six male subjects (average age 34 years) attending the Nanjing (China) STD Clinic, who presented with urethral discharge and/or dysuria and were diagnosed with *N. gonorrhoeae* infection (**Table 1**). Four men were co-infected with *Chlamydia trachomatis* and/or *Mycoplasma genitalium*; three had a history of prior gonococcal infection and three had self-administered antibiotics prior to their clinic visit (**Table 1**). In contrast, none of the seven enrolled women had a prior history of gonococcal infection or had taken antibiotics prior to their clinic visit (**Table S2**). Two gonococcal strains were isolated from a man and a woman who comprised a single dyad (subjects M-4 and F-6).

**Table 1.**
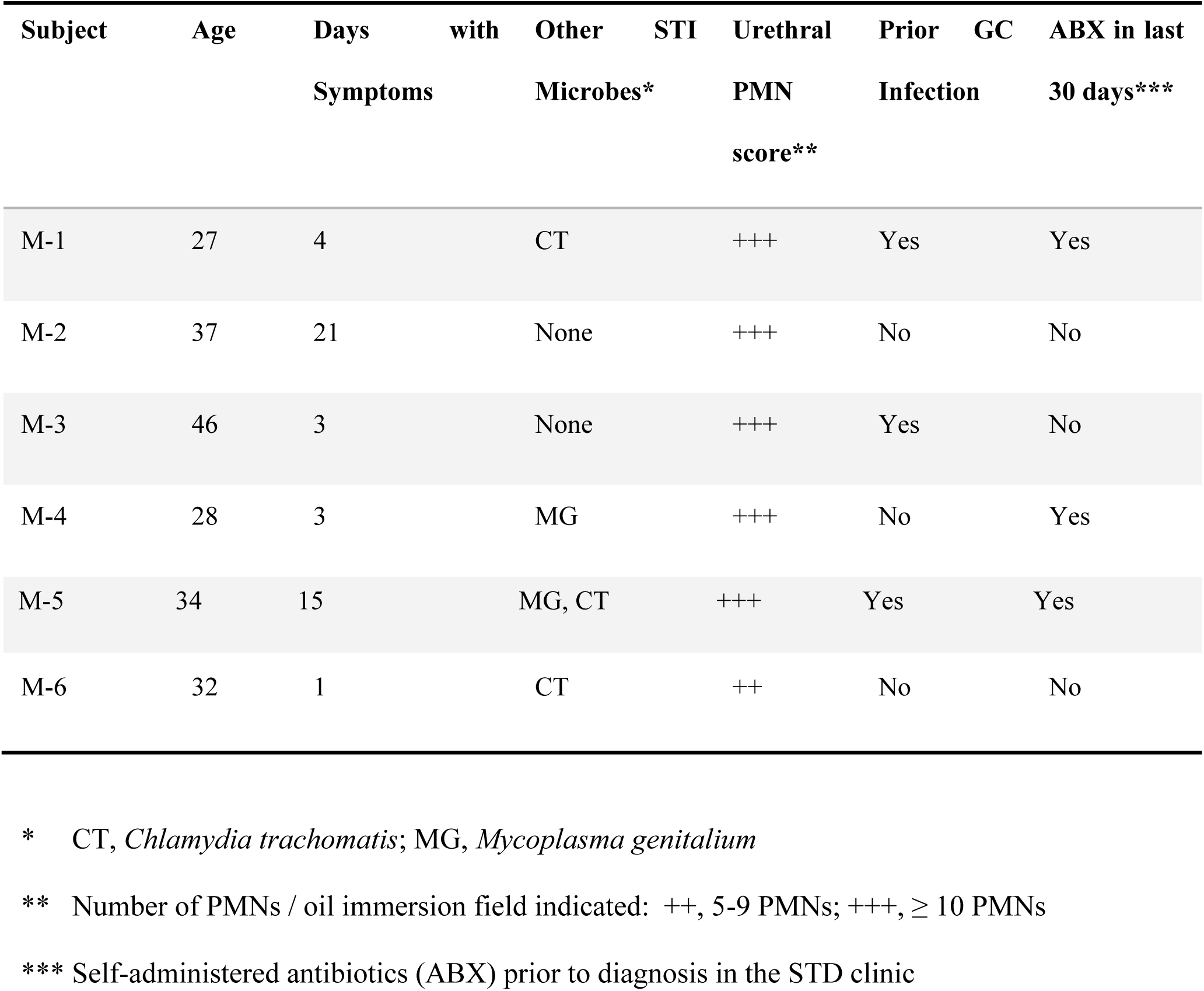
Characteristics of male subjects with gonococcal urethritis

### Gonococcal Gene Expression in the Human Genital Tract

RNA was extracted from male urethral or cervico-vaginal lavage specimens and individual cDNA reads aligned to *N. gonorrhoeae* strain FA1090 using Rockhopper (39). Total gene expression was reported in reads per kilobase per million (RPKM). On average, 8.1% of the total RNA from male urethral specimens, aligned to the *N. gonorrhoeae* FA1090 genome and 42.5% to the human genome; 4.3% of total RNA from cervico-vaginal lavages aligned to FA1090 and 45% to the human genome.

Expression of 1725 gonococcal genes (approximately 65% of the gonococcal genome) was detected in male urethral specimens. High expression of a core set of genes that encoded for oxidative stress products (eg. peroxiredoxin/glutaredoxin), housekeeping genes (eg. elongation factor Tu), outer membrane proteins (eg. Rmp [Omp3] and porin P.IB) and hypothetical proteins were identified across male specimens (**Table S3**). A more detailed analysis of the top 200 gonococcal genes (approximately 10% of total genome) that were expressed in all six male urethral secretions (in the aggregate) was carried out by determining the ratio of gene enrichment and segregation into broad categories. Despite some variability in RPKM values among subjects, significantly enriched categories included pilin, oxidative stress and regulatory proteins (**Figure 1**), which reflected a common pattern of high gene expression during natural infection.

**Figure 1.**
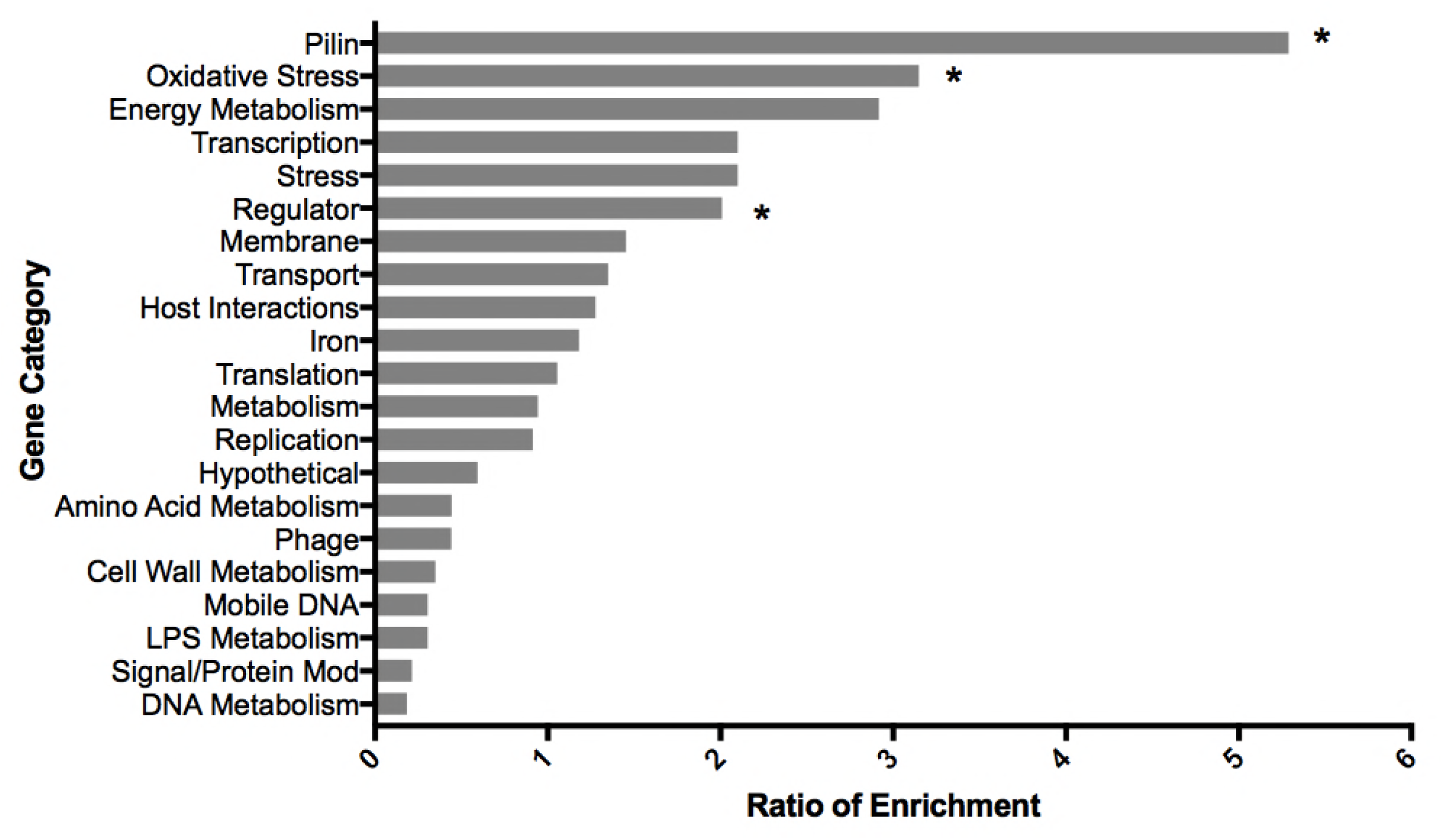
Categorization of *N. gonorrhoeae* top 200 genes expressed during natural infection in men. Expression of gonococcal genes in 6 infected men was averaged and the top 200 genes were categorized based on characteristics indicated for *N. gonorrhoeae* strain FA1090 available in NCBI. Categories are shown on the y-axis and the ratio of enrichment (percentage of genes of a given functional category in the RNA-seq dataset / percentage of genes assigned to that functional category in the whole gonococcal genome) on the x-axis. For clarity, ribosomal protein, tRNA, and rRNA genes were removed. Asterisks (*) indicate that enrichment was increased significantly (p-value ≤ 0.5); Fisher’s exact test.

### Gonococcal Gene Regulation in the Male Genital Tract

Our earlier studies had demonstrated that a large subset of gonococcal genes was regulated *in vivo* during cervical infection in women (38). Gonococcal gene expression in the male genital tract was compared to expression of matched infecting strains grown *in vitro* by analysis of RNA seq data sets for individual strains and for the composite datasets. Genes that displayed a significant difference in expression during *in vivo* vs. *in vitro* conditions were selected for further analysis based on the following criteria: a ≥ 2-fold difference in RPKM value, a q value ≤ 0.05 and expression greater than 10 RPKM under at least one condition (**Table S4**). The majority of genes were expressed similarly *in vivo* and *in vitro*, but approximately 4.2% of the total number of genes detected (79 genes) revealed increased expression *in vivo* compared to that observed *in vitro*; expression of 5.2% (99 genes) was decreased *in vivo* compared to that observed *in vitro* (**Figure 2A**).

**Figure 2.**
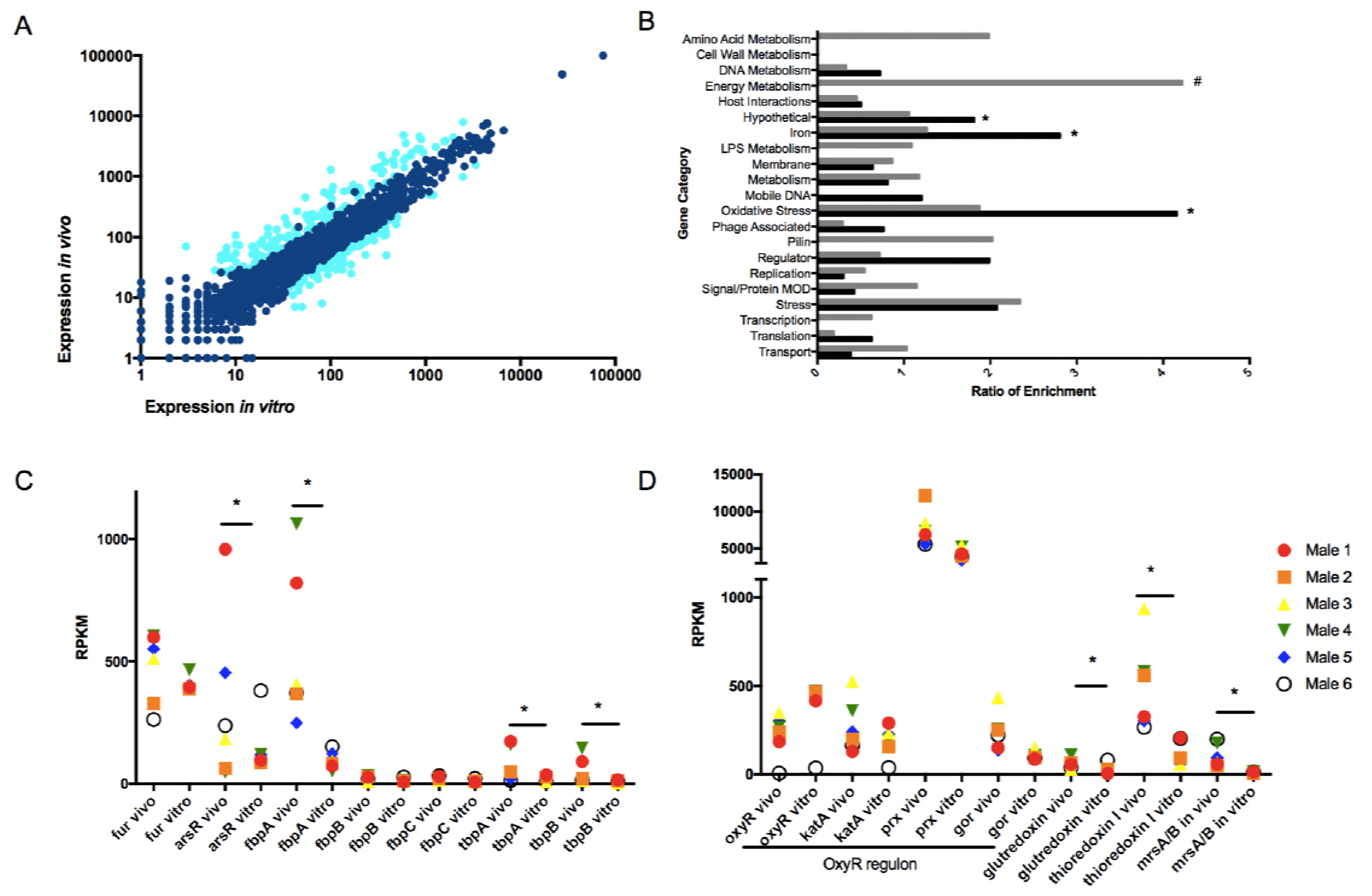
Comparison of *N. gonorrhoeae* gene expression in infected men, *in vivo*, and isolates grown *in vitro*. (**A**) Composite gene expression levels in urethral specimens (n=6) on the y-axis plotted against composite expression of corresponding gonococcal isolates (n=6) grown *in vitro* (x-axis). Shown as log_10_ expression (RPKM) levels. Genes indicated in light blue (upper genes) had q-values ≤ 0.05 and ≥ 2-fold changes in *in vivo* (vs. *in* vitro) expression and genes in light blue (lower genes) had q-values ≤ 0.05 and ≤ 2-fold changes in *in vivo* expression. (**B**) Functional enrichment of genes with statistically significant changes in expression under *in vivo* and *in vitro* conditions. Genes were differentially expressed if the q value was ≤ 0.05 and the fold change ≥ 2. Grey bars: functional enrichment of genes with increased expression *in vivo*. White bars: functional enrichment of genes with decreased expression *in vivo*. Categories are shown on the y-axis and the ratio of enrichment (percentage of genes of a given functional category in the RNA-seq dataset / percentage of genes assigned to that functional category in the whole gonococcal genome) on the x-axis. Ribosomal protein, tRNA and rRNA genes were removed for clarity. * indicate significant enrichment amongst genes increased *in vivo* (vs. *in vitro)* and # indicate significant enrichment amongst genes decreased *in vivo*, with a p-value of ≤ 0.05 by Fisher’s exact test. (**C**) Expression levels of iron genes and (**D**) oxidative stress genes under *in vivo* and *in vitro* conditions (6 male specimens [in vivo] and the corresponding 6 isolates [in *vitro]* are shown in the same color in adjacent columns). * Indicates a q-value of ≤ 0.05.

Regulation of gonococcal gene expression was then examined by functional enrichment analysis (**Figure 2B**). Genes involved in iron and oxidative stress pathways and genes encoding hypothetical proteins were among those significantly enriched and were more highly expressed *in vivo* than *in vitro*. For example, expression of certain iron scavenging genes, eg. *fbpA, tbpA* and *tbpB* and of the *fur* controlled regulator *arsR* was significantly higher *in vivo* than *in vitro*, although variance in *arsR in vivo* expression was large (**Figure 2C**). This observation suggests that the male genital tract is iron deplete (similar to the female genital tract (38)). Genes involved in energy metabolism were also significantly enriched but had decreased expression *in vivo* (**Figure 2**). Among the down-regulated energy metabolism genes was the *nuoA* / *nuoN* operon (NGO1737-1751) (40), involved in aerobic respiration; its overall decreased expression *in vivo* may suggest a mechanism for bacterial survival in an anaerobic environment (41).

Among the enriched oxidative stress genes increased *in vivo*, we observed a significant increase in a glutaredoxin family protein (NGO0031), *thioredoxin I trx1* (NGO0652) and *mrsAB* (NGO2059) (**Figure 2D**). The family of glutaredoxin proteins function as glutathione-dependent reductases (42); *mrsAB* plays a role in reducing oxidized methionines to reactive peptides (43) and *trx1* has been shown to respond to oxidation and is under control of the Fur regulon (44, 45). While expression of the well-studied OxyR regulon *(oxyR, prx, kat*, and *gor)* was not significantly different in *in vivo* samples compared to cultures grown *in vitro*, we observed decreased expression of the OxyR repressor *in vivo*, with increased expression of genes within the OxyR regulon *(prx, kat*, and *gor)*. This notable trend further suggests that the gonococcus is exposed to reactive oxygen species during mucosal infection in men (**Figure 2D**) (46, 47).

Several gonococcal non-coding RNAs were also regulated during infection in men. These included 4 rRNAs and 29 tRNAs, which were expressed at higher levels during infection compared to growth *in vitro* except for NGO t31, an arginine tRNA, that was expressed more highly during *in vitro* growth (**Table S5**). Putative sRNAs (defined as expression representing an intergenic region of the complementary strand of a known ORF and 30–250 nucleotides in length) were also regulated during infection; 29 sRNAs were increased *in vivo* compared to *in vitro* and 21 sRNAs were decreased *in vivo* (**Table S6**). Among these sRNAs, several have been described previously (48, 49).

### Comparison of Gonococcal Gene Signatures in the Male and Female Genital Tract

Building on gonococcal transcriptome data in infected men and women, we examined whether site-specific, environmental conditions influenced expression of gonococcal genes during infection. A comparative transcriptome analysis was carried out using the 6 urethral and 7 cervico-vaginal specimens. We performed principle component analysis (PCA) to assess whether global *in vivo* gene expression revealed transcriptional signatures specific for infections in men or women (**Figure 3A**). PCA showed a clear distinction between gonococcal genes expressed in specimens from men and women, with high variance among the 7 specimens from women compared to low variance in men, as evidenced by tight clustering when the first two principal components were plotted against each other (**Figure 3A**). Gene expression analysis of the composite datasets was also performed; parameters set for defining differential gene regulation were: a ≥ 2-fold change in RPKM, a q value ≤ 0.05 and a difference in expression level of at least 10 RPKM between men and women. Of 1879 genes expressed, 176 (9.4%) were increased in men compared to women and 80 (4.3%) were decreased in men indicating that nearly 14% of gonococcal genes were differentially regulated during gonococcal infection in the male and female genital tracts (**Table S7**).

**Figure 3.**
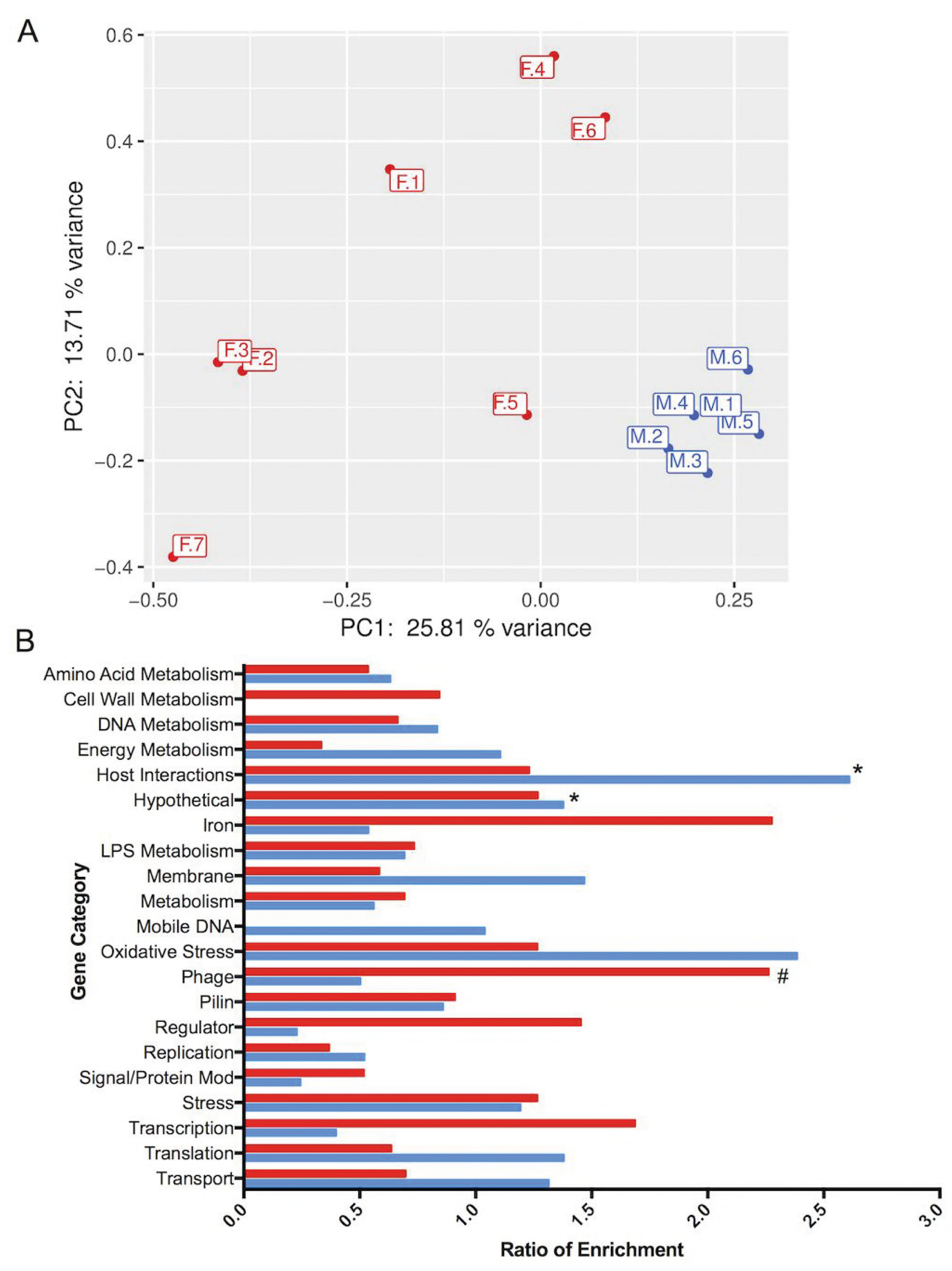
Comparison of *N. gonorrhoeae* gene expression during infection in men and women, *in vivo*. (**A**) Variance in global expression in each specimen from men and women. The first principal component (PC1, x axis) had a variance of ~26% and the second principal component (PC2, y axis) had a variance of ~14%. (**B**) Functional enrichment of genes with significant changes in expression levels in men and women. Genes were differentially expressed if the q value was ≤ 0.05 and the fold change was ≥ 2. Blue bars: genes with increased expression, *in vivo*, in men compared to women. Red bars: genes with increased expression, *in vivo*, in women compared to men.

A functional enrichment analysis of gene categories performed on male and female specimens revealed that phage associated genes were significantly enriched in female specimens (**Figure 3B**). While female specimens had higher expression of a broad range of phage types (eg. double stranded and filamentous single stranded phages), male specimens had higher expression in the subset of double stranded DNA prophages (**Figure S1A**) (50). Other highly expressed categories enriched in male specimens included genes that encoded hypothetical proteins and proteins involved in host interactions (**Figure 3B**). Iron and oxidative stress genes that were differentially expressed in male and female specimens included: ferredoxin (NGO1859); a TonB receptor protein (NGO1205) and *fetA* (NGO2093; *fetA* is regulated by MpeR and induced in low iron conditions (51)). These genes all manifested increased expression in women (**Figure S1B**). In contrast, expression of the oxidative stress gene *bfrB* was increased in male specimens; other oxidative stress genes were similarly expressed in both men and women (**Figure S1C**), suggesting that exposure to oxidative stress occurs in both the male and female genital tracts.

Regulation of several gonococcal sRNAs was also observed. A significant change in expression of 119 sRNAs was observed in specimens obtained from men and women; 28 were expressed at higher levels in male specimens and 91 in female specimens. Previously identified sRNAs showing regulation based on sex included smRNA4, smRNA5, smRNA8 (49), *fnrS* (a small RNA induced under anaerobic conditions (44)) and the iron-repressed sRNA encoding NrrF (52) (**Table S6**). In addition, 45 tRNAs showed differential regulation with all being more highly expressed in female patients compared to male patients (**Table S5**).

### Expression of Antimicrobial Resistance Genes

Gonococcal antimicrobial resistance (AMR) is a growing concern and is common in strains from the eastern regions of the world, including China. We examined phenotypic and genotypic AMR patterns from our cohort and found that all the isolates exhibited resistance to penicillin, tetracycline, and ciprofloxacin; one isolate was resistant to azithromycin (**Table 2**). One isolate also showed reduced susceptibility to ceftriaxone (**Table 2**). Strains isolated from women exhibited similar antimicrobial susceptibility patterns (**Table 2**). These results were similar to AMR patterns of male urethritis strains reported earlier (2011–12) from the Nanjing STD clinic (53).

**Table 2.**
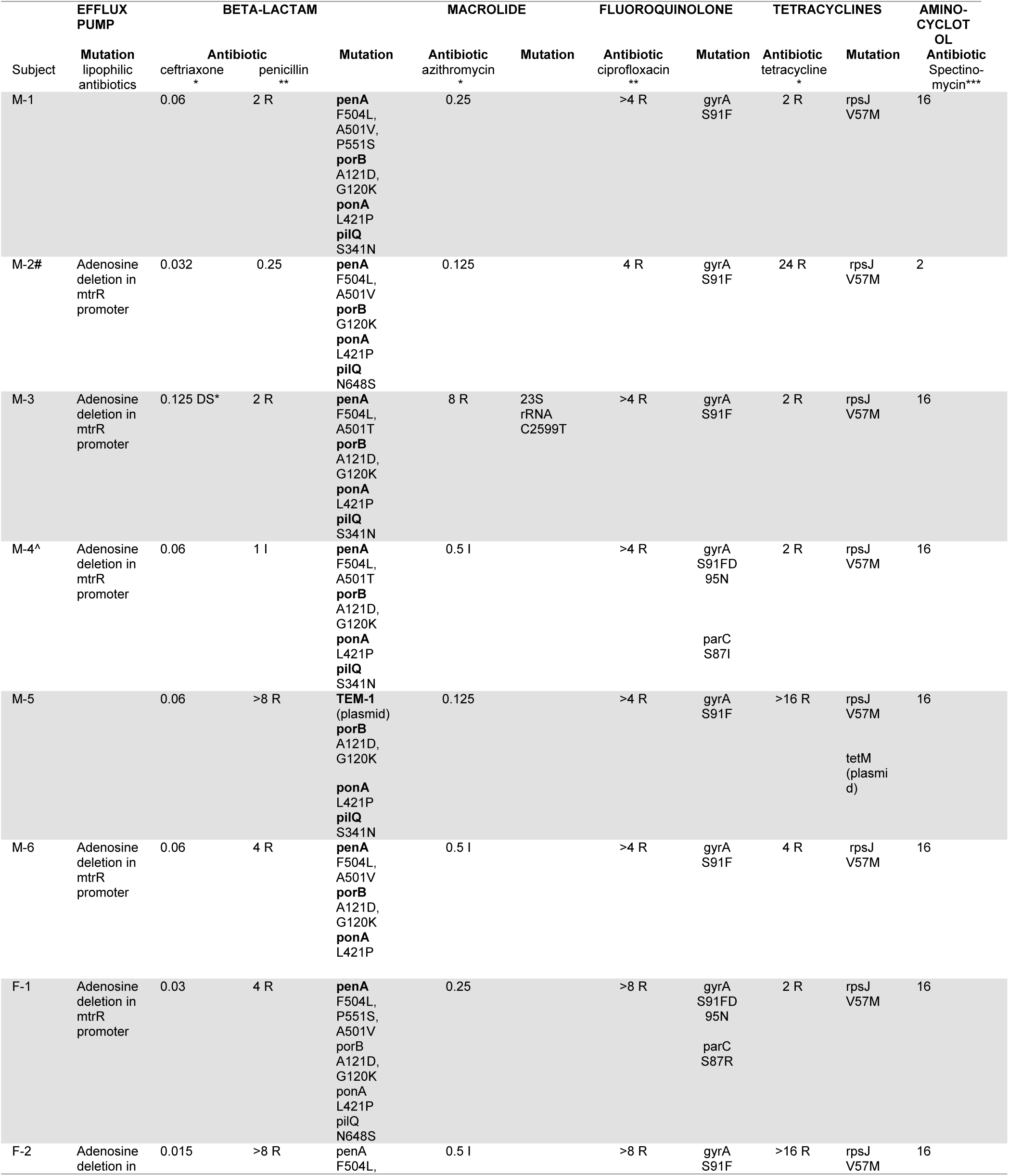
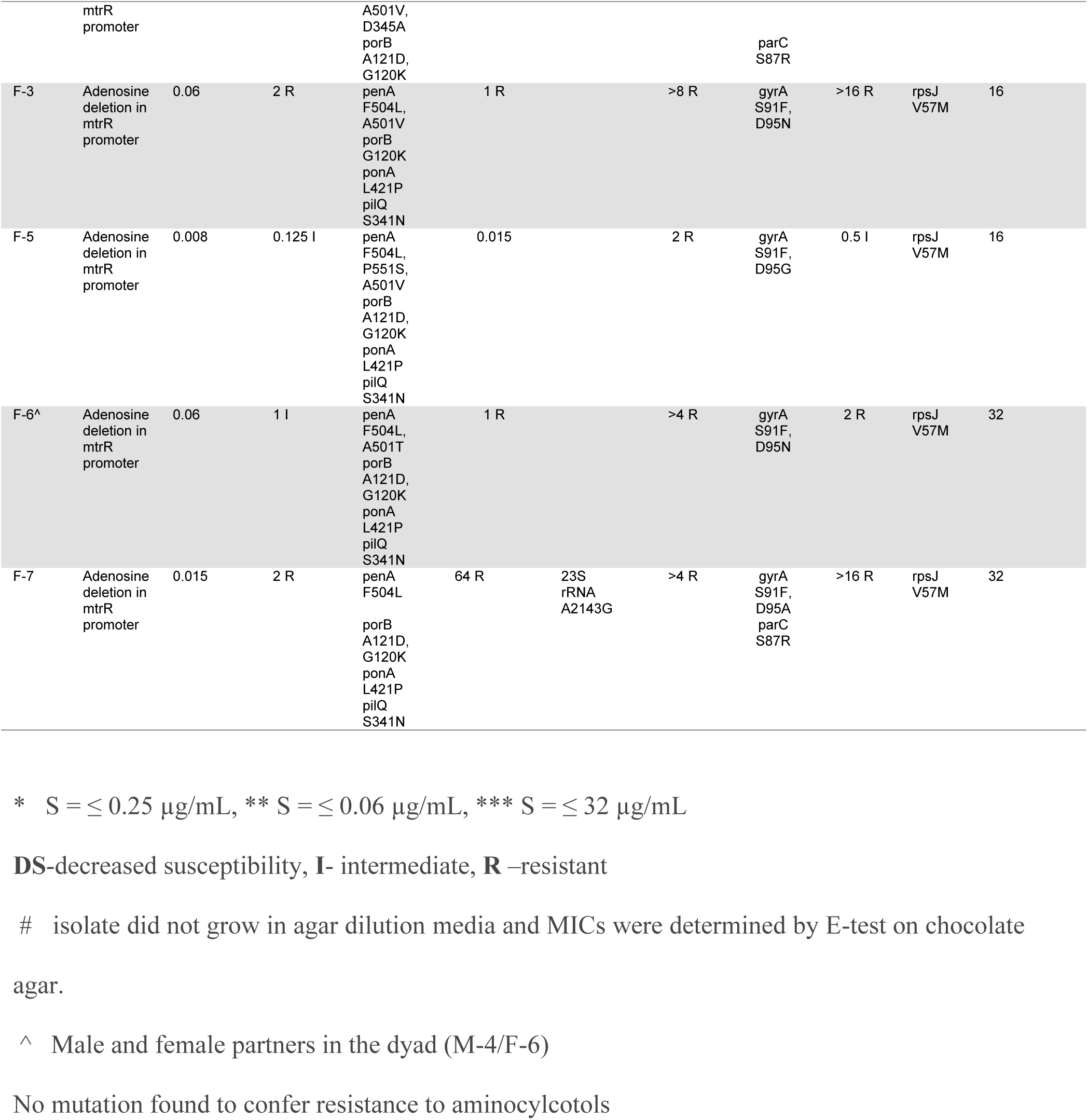
Phenotypic resistance and AMR genes identified in male and female *N. gonorrhoeae* isolates (n=13) by WGS

Whole genome sequencing was performed on all available strains (6 male and 6 female) to define strain relatedness and genotypic determinants of antibiotic resistance. A single nucleotide polymorphism (SNP) distance tree was constructed that used SNP distances (differences) that ranged between 81 and 3913 separating the genomes (**Figure S2**). The male and female dyad (M-4 and F-6) yielded genomic sequences that were separated by only 81 SNPs. Strain NCCP11945 isolated from a subject with gonococcal disease in Korea (average SNP distance of 2828 SNPs compared to strain FA1090) shared considerable homology with the isolates obtained from our cohort as compared to the commonly used laboratory strain, FA1090 (average distance of 4212 compared to FA1090).

Genomic analyses based on publicly available, curated, antibiotic resistance databases (CARD and GC-MLST), identified multiple AMR genes (**Table 2**) and SNP-level variants that likely contributed to phenotypic resistance. All gonococcal strains exhibited SNP mutations in *gyrA* and/or *parC* (which confer resistance to fluoroquinolones), SNP mutations in *penA, ponA, porB*, and *pilQ* (beta-lactams resistance), and SNP mutations in *rpsJ* (tetracycline resistance) (54). Only one isolate carried plasmid-based resistance genes (M-5: *TEM-1* and *tetM)*. Interestingly, among the male subjects, M-3 was the only isolate to exhibit azithromycin resistance and a corresponding mutation in the 23s rRNA gene (55). Three isolates had an adenosine deletion in the promoter region of *mtrR*, the repressor of the *mtrCDE* efflux pump, a system that exports bactericidal host-derived compounds and hydrophobic antibiotics (beta-lactams and macrolides) from the bacterial cell (56–58). Isolates from women exhibited similar genomic patterns of resistance: by and large the same mutations identified in the male isolates were also present in the female isolates (**Table 2**). Among 6 SNPs associated with beta-lactam resistance in the dyad isolates (subjects M-4 and F-6 who were infected by the same strain), 5 SNPs were shared in the dyad isolates. Two SNPs in *gyrA* were shared (S91F and D95N) and a Ser SNP in *parC* (S87I) was detected only in the male isolate. Finally, while several SNP mutations coincided with phenotypic resistance to azithromycin, ciprofloxacin and penicillin (**Table 2**), the single isolate with decreased sensitivity to ceftriaxone (M-3) did not possess the mosaic *penA* XXXIV allele (59), nor an additional curated mutation(s) that was absent in the phenotypically sensitive strains (**Table 2**) (59, 60).

Expression levels of genes associated with antibiotic resistance in men were also compared to expression of corresponding genes *in vitro* (**Figure 4A** and B). Expression of the *pilQ* gene, associated with antibiotic permeability (14), was significantly decreased by 1.78-fold *in vivo* compared to *in vitro* (**Figure 4A**). The MisRS two-component regulatory system is important for resistance to host-derived antimicrobial peptides and antibiotics (61); and though not statistically significant, *misR* had 2.08-fold higher expression *in vivo* than *in vitro*, supporting its important role *in vivo*. The expression of the Mtr efflux pump was also closely examined; expression of *mtrA*, which can influence expression of *mtrCDE* (62), was significantly decreased by 2.75 fold in *vivo* (**Figure 4B**). Gonococcal strains from male urethral specimens (M-2, M-3, M-4, M-6) contained an adenosine deletion in the *mtrR* promoter that was accompanied by the lowest expression of *mtrR* and highest expression of *mtrCDE* and *mtrF*, confirming that the adenosine deletion directly affects expression of the Mtr complex (63) *in vivo*. Expression of *mpeR* was also investigated, as it has been shown to regulate the Mtr locus, however, expression was below 10 RPKM in all *in vitro* and *in vivo* samples.

**Figure 4.**
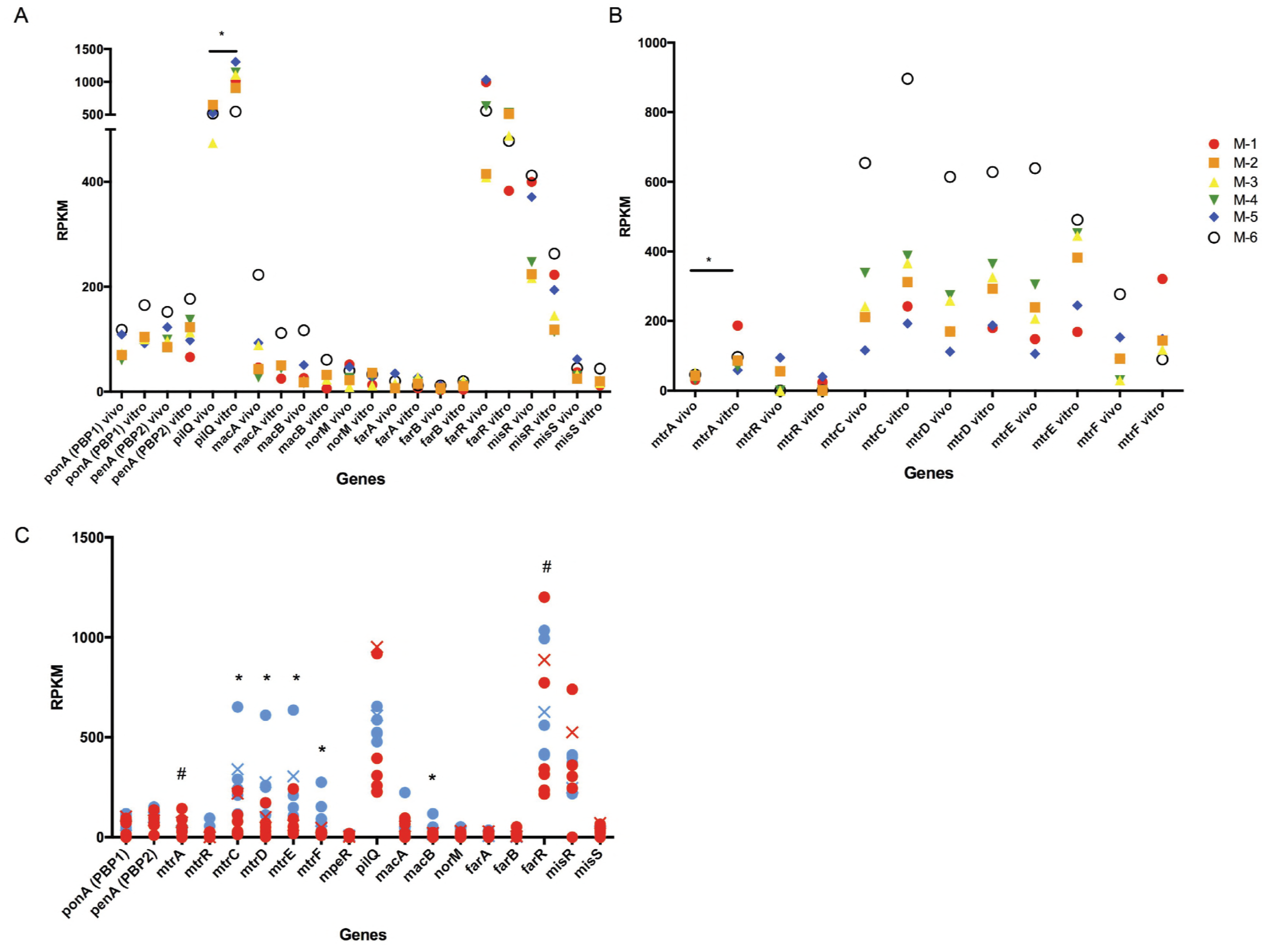
Expression of gonococcal antibiotic resistance genes. Expression levels of antibiotic resistance genes. (**A**) and the Mtr efflux pump (**B**) in the male genital tract (*in vivo*) and the corresponding strain in CDM *(in vitro);* genes identified on the x-axis (6 male specimens [in vivo] and the corresponding 6 isolates [in *vitro]* are shown in the same color). *Indicates a q-value of ≤ 0.05. (**C**) Expression levels of antibiotic resistance genes from the male genital tract (blue) or the female genital tract (red). * Indicates higher genes expression in men and # higher in women (q-value of ≤ 0.05). Xs indicate the male (M-4) and female (F-6) counterparts of the dyad.

Transcriptomes from male urethral specimens were also compared with female cervico-vaginal lavages to determine if expression of AMR genes was influenced by sex (**Figure 4C**). Because of the adenosine deletion in the *mtrR* promoter in 4/6 males (see above) and all female isolates (**Table 2**), expression of *mtrR* was low in both groups but resulted in significantly higher expression of *mtrCDE* and *mtrF* in men, as predicted (63), but not in women. Even within the dyad where both M-4 and F-6 contained the adenosine deletion in the *mtrR* promoter, M-4 had approximately 2x higher expression levels of *mtrCDE* than F-6. Interestingly, the expression of *mtrA*, an activator of *mtrCDE* (62), was significantly higher during infection in women as compared to men, despite lower expression of *mtrCDE* in women (**Figure 4C**). The *farR* gene, which negatively controls expression of the *farAB* operon by directly binding to the *farAB* promoter (64) to prevent excess expression of gonococcal efflux pumps, also showed significantly higher expression in cervico-vaginal lavages. Collectively these results suggest that, although strains isolated from infected men and women exhibit comparable AMR genotypes and *in vitro* phenotypes, expression of AMR genes is different during infection in the male and female genital tract.

## Discussion

*N. gonorrhoeae* infects the male and female genital tract, two very distinct environments in humans. As there is little doubt about the intrinsic tissue, cellular and molecular differences that define these host environments, it is reasonable to assume that the gonococcus will adapt to these environmental differences during infection (65). Building on our previous results on the gonococcal transcriptome profile during infection in women (37), we extended our analysis to specimens from infected men. In the present study, we not only report a distinct gonococcal transcriptome profile *in vivo* as compared to *in vitro* in infected men, but also, for the first time, describe a comparison of *N. gonorrhoeae* gene expression during infection in the male and female genital tract. Our results also demonstrate distinct gonococcal gene expression signatures in men and women, consistent with the intrinsically different make-up and nature of the two sites of infection. Furthermore, our approach based on both whole genome- and RNA-sequencing, enabled us to evaluate expression of antibiotic resistance determinants during infection in both the male and female genital tract.

Currently, the only human model of gonococcal infection is experimental urethral challenge of male volunteers. Those studies have examined gonococcal virulence factors and have provided valuable information on opacity proteins, genes involved in iron acquisition, structure modification of lipooligosacharides and a variety of other established virulence factors (32, 66, 67) but are limited by the short duration of experimental infection. The small number of infecting strains used experimentally in men (mostly laboratory-adapted strains FA1090 and MS11mkC (32) have precluded an in-depth understanding of the range of adaptions employed during gonococcal infections, generally. Using a panel of *N. gonorrhoeae* strains isolated from infected subjects, we compared the gonococcal *in vivo* transcriptome expressed during infection of the male genital tract to that of the corresponding infecting strains grown *in vitro*. Our results revealed increased expression of genes involved in oxidative stress and iron scavenging *in vivo*, accompanied by decreased expression of genes involved in metabolism-associated categories (i.e. energy and amino acid metabolism). In particular, we observed that gonococci expressed high levels of *trx1, msrAB* and glutaredoxin genes during infection in male subjects, in agreement with the large PMN influx that occurs during infection in symptomatic men. Our results also demonstrated increased expression of iron-regulated genes *(tbpAB* (66)), indicating that the male genital tract is an iron-deplete environment. While we recognize that *in vitro* culture conditions differ compared to the conditions during the human genital tract beyond metabolite content, this study provides a gene specific view of the response of *N. gonorrheoae* to infection and leads to a number of hypotheses regarding infection mechanisms that can be explored in more detail in future studies.

While we identified both expected and novel differences in gonococcal gene expression during human infection compared to *in vitro* growth, the most revealing aspect of our study was the differential gonococcal expression profile observed in men and women. Our results showed that nearly 14% of gonococcal genes that were expressed in human infection were differentially expressed in men and women. Principle component analysis demonstrated that gonococcal expression in men and women separated into distinct gene expression signatures, with high variance among the 7 female specimens and low variance among the 6 male specimens. We acknowledge that high variance could be due to the small sample size, but may also be due to other female-specific environmental factors that could alter gonococcal gene expression, such as menstrual cycle or the microbiome, which exhibits diversity among individuals (68). When genomic DNA sequence analysis was also considered, there was no clustering among strains derived from men or women (except for strains in the single dyad), which strongly suggests that the observed differences in transcriptomic signatures were driven by exposure of gonococci, generally, to the male or the female genital tract and not by strain relatedness. Furthermore, this is also supported in the single dyad by differences in gene expression manifested by the same strain in each partner, which reflected changes in gene expression that were seen overall in the other men and women. In addition to the small sample size, it is important to also consider other limitations of the study when interpreting sex-specific and growth-specific aspects of *N. gonorrhoeae* infection. Due to the design of the study, men are recruited because they enter the clinic with a symptomatic response, thus, men may have a similar stage of infection and gonococcal growth stage; this is in contrast to the matched females, who are partners of the recruited men and who were at various stages of infection and therefore gonococcal growth.

Among the most relevant gonococcal transcriptome differences that emerged in male and female specimens, were in genes encoding anti-microbial resistance determinants. The rise of antibiotic resistance in *N. gonorrhoeae* is now a global concern (69). Patterns of resistance vary among countries because of population mobility and geographically-localized increases in antimicrobial resistance--such as those observed in China (70); these patterns are predictive of future resistance patterns in countries, which currently have lower levels of resistance, such as the US (71, 72). Whole genome DNA sequencing (WGS) of the gonococcal strains isolated from our male and female cohort revealed similarity of genetic determinants of resistance to strains isolated in the US (59). In all strains, regardless of the origin of the specimens (men or women), the presence of an adenosine deletion in the *mtrR* promoter (also detected by WGS) correlated with increased expression of the Mtr regulon. Nevertheless, strains isolated from men showed a significantly higher expression of *mtrCDE* and *mtrF* than those from women. While a direct explanation for these results remains speculative, it is possible that biofilm formation in the female genital tract (and lack of it in the male genital tract) is a contributing factor (19, 20). Biofilms can provide protection against harsh environments and their absence in the male genital tract may lead to increased expression of antibiotic resistant efflux pumps, as a defense mechanism (73). An additional explanation for higher expression of *mtrCDE* in men have may have been the ready accessibility of antibiotics in China; approximately half of the male subjects (but none of the female subjects) in this study had self-administered antibiotics prior to seeking care at the STD clinic. Future studies including a larger cohort may help resolve these differences.

*N. gonorrhoeae* contains phage DNA inserted in the genome; five regions comprising dsDNA lysogenic phage genomes (50) and four additional regions comprising filamentous phages (74). The role of phage genes in *N. gonorrhoeae* infection has not been studied extensively. Previous work from our group has identified a phage gene, *npr*, involved in invasion of female epithelial cells *in vitro*, which correlates with disease progression in a female mouse model of gonococcal infection (75). Our finding that expression of phage genes was increased in female subjects may also relate to the formation of biofilms in the female genital tract (20). We propose that biofilm formation may facilitate DNA exchange via phage among bacteria, thus contributing to differences in infections of men and women. Additional transcriptional patterns that differentiated organisms obtained during infection in men and women included genes encoding tRNAs with higher expression observed in women as compared to men. However, apparent expression of tRNAs may be partially affected by reads aligning to additional microbes present in the genital tract: tRNA genes are highly conserved across species and the women in the study may have a larger population of microbial organisms. Six of seven gonococcal infected women reported in this study had BV and/or other bacterial genital infections. Some studies have suggested a role for tRNAs in bacterial infection (76, 77), and it is possible that differences observed in men and women also correlate with the enhanced microbial environment in women. Small RNA expression *in vivo* was also different; NrrF and FnrS (an sRNA induced in anaerobic conditions) were expressed at higher levels in women than in men.

Collectively, our results provide the first global view of gonococcal gene expression during infection in humans and define gene signatures specific to infections in men and women. We report important differences related to antibiotic resistance and gonococcal pathogenesis that can be extrapolated to improve understanding gonococcal disease outcomes and potential treatments. Our analysis also highlights shortfalls of studying bacterial infections using *in vitro* models and systems. It is critical that studies designed to identify targeted therapies for gonococcal infections consider sex-specific differences in gene expression profiles that may impact treatment outcomes. Furthermore, addressing how expression of antimicrobial resistance genes are driven by environmental cues in the male and female genital tract has important implications for the use of targeted antibiotics.

## Methods

### Ethics statement

All subjects provided written informed consent in accordance with requirements by Institutional Review Boards from: Tufts University, Boston MA (protocol number 11710); the University of Massachusetts Medical School, Worcester, MA and Boston Univeristy School of Medicine, Boston MA; the Institute of Dermatology, Chinese Academy of Medical Sciences & Peking Union Medical College, Nanjing, China.

### Identification of subjects and characterization of urethral infections

Urethral swab specimens were collected from six men attending the Nanjing Sexually Transmitted Disease (STD) Clinic, chosen at random, who were participants in a study of gonococcal transmission (78). Male subjects presented with symptoms and signs of urethritis (dysuria and/or urethral discharge). Gram stains of urethral exudates showed PMNs and Gram-negative intracellular diplococci, criteria that are highly specific (99%) for gonococcal infection (79). Urethral swab specimens from men with urethritis were also inoculated onto Thayer-Martin medium (DL Biotech, China) and cultured in candle jars at 36°C. Gonococcal isolates were identified by colonial morphology, Gram’s stain, and oxidase testing. Urethral swab specimens were also tested for *Chlamydia trachomatis* and *Mycoplasma genitalium* by PCR (80, 81), and *Ureaplasma urealyticum* and *Mycoplasma hominis* by culture using Mycoplasma IST2 test (bioMerieux, France).

Collection of cervico-vaginal lavage specimens (n=4) was described previously (38) and three additional specimens were obtained for the present study (n_to_t_al_ =7). PCR testing using primers directed against the gonococcal *porA* pseudogene (82) had previously identified *N. gonorrhoeae* infection in one woman (F-4) whose cervical culture was negative. Women reported that they were monogamous (3/7 were married), having been the sole sex-partner of a man with gonococcal urethritis. One woman (F-6) and one man (M-4) were members of a single dyad.

### Antimicrobial testing of N. gonorrhoeae

*N. gonorrhoeae* strains from urethral swab and cervico-vaginal lavage specimens were identified and processed as previously described (38). Antimicrobial sensitivity testing of *N. gonorrhoeae* isolates was determined for penicillin, tetracycline, spectinomycin, azithromycin, ceftriaxone, cefixime and ciprofloxacin at the Nanjing STD Clinic. Mean inhibitory concentration (MICs) were determined by the agar dilution method and current Clinical and Laboratory Standards Institute cutoffs were used, except for azithromycin, in which we followed the European Committee on Antimicrobial Susceptibility Testing; www.eucast.org (83). A single isolate (from M-2) that did not grow in agar dilution media, nor GC agar, had MICs determined on chocolate agar by E-test, thus MIC values may be one dilution lower due to enriched agar (84).

### RNA isolation

Total RNA was isolated from urethral swab and cervico-vaginal lavage specimens using TRIzol (Invitrogen) as described (38). Briefly, specimens were washed twice with 70% ethanol and treated with TURBO DNase (Ambion), RiboMinus Kit (Invitrogen) (to deplete eukaryotic rRNA) and Microbe EXPRESS Kit (Ambion) (to deplete bacterial rRNA) using diethylpyrocarbonate (DEPC)-treated EDTA-free water. Gonococcal isolates that corresponded to each specimen were grown overnight on chocolate agar plates at 37°C in 5% CO_2_ prior to inoculation in chemically defined media (CDM) containing ferric nitrate (100 μM). Liquid cultures were inoculated at an O.D_600_ of 0.1, harvested after 3 hr., pelleted and RNA extracted as described above.

### RNA sequencing and analysis

RNA sequencing was performed as previously described (38). cDNA libraries were prepared with TruSeq RNA Preparation Kit and sequenced on an Illumina HiSeq 2500 instrument using High Output V3 chemistry in single read 100 formats. Analysis of RNA-seq. data was performed using the Rockhopper program, aligning the reads to the NCBI genome sequence of *N. gonorrhoeae* strain FA1090, and then reporting RNA-seq. data in reads per kilobase per million (RPKM) (37). The cut-off for gene expression was set at an RPKM value of at least 10. A composite set of RPKM values was derived from RNA-seq. data from the six male urethral and another for the three newly collected cervico-vaginal added to the four reported earlier (Gene Expression Omnibus database with accession number GSE71151) (38). Henceforth, these are referred to as “*in vivo";* RNA-seq. Data from gonococcal strains isolated from each subject and cultured *in vitro* were also obtained; referred to as “*in vitro"*.

Statistical analysis was performed on pooled data from each category of specimens: *in vivo* urethral; *in vivo* cervico–vaginal lavage and *in vitro*. Genes with greater than a 2-fold change and a *q*-value of ≤ 0.05 in each category were deemed as regulated. Gene assignments and category were performed using KEGG and NCBI RefSeq public databases. Functional enrichment ratios were determined by dividing the percentage of genes in a functional category for a particular set of genes (e.g. highly expressed genes, differentially regulated genes) by the percentage of genes in the same category present in the gonococcal genome as a whole. This ratio was used to interpret enrichment of a particular set of genes that possessed a specific function. Fisher’s exact test was used to assign a p-value to the enrichment. Principal Component Analysis (PCA) (85) was performed in R using singular value decomposition on the scaled and centered data set of RPKM values for all genes in each subject, and the resulting data visualized by sex on a colored plot.

### Whole-genome sequencing and analysis

*N. gonorrhoeae* isolates were grown on GC agar plates supplemented with 1% IsoVitaleX™ and 0.6% fetal bovine serum at 37°C and 5% CO_2_ for 16–18 hr. Genomic DNA was extracted from a 10 μL inoculum of each bacterial culture using a 5PRIME DNA extraction kit (5Prime, San Francisco, CA), following the manufacturer’s recommendations with slight modifications. Whole-genome sequencing (WGS) was conducted using a PacBio RSII platform (Pacific Biosciences, Menlo Park, CA) with P5-C3 chemistry. *De novo* genome assembly was conducted using the hierarchical genome assembly process workflow (HGAP3, SMRTAnalysis 2.3.0), which included consensus polishing using Quiver (86). Single-nucleotide polymorphisms (SNPs) across sequenced isolates were called *de* novo-assembled contigs using kSNP3 without a reference strain (87). Concatenated SNPs were used to construct phylogenetic trees using the approximate-maximum-likelihood-based approach; trees were visualized in FigTree (http://tree.bio.ed.ac.uk/software/figtree/). Antibiotic resistance elements were identified using the Comprehensive Antibiotic Resistance Database (88) and the PubMLST Neisseria database (https://pubmlst.org/neisseria) (89).

### Data Availability

New RNA-seq. data from this study has been deposited in the Gene Expression Omnibus (GSE113290). Genomic data from isolates has been deposited into NCBI under project number PRJNA329501 (**Table S1**).

## Acknowledgements

This work was supported by NIH/NIAID grants U19AI084048 (Rice and Genco) and RO1AI048611 (Genco). PNNL is operated for the DOE by Battelle Memorial Institute under contract DE-AC05-76RLO 1830. The conclusions, findings and opinions expressed by the authors do not necessarily reflect the official position of the US Centers for Disease Control and Prevention (CDC).

---

**Supplemental Figure 1.** Phage expression, iron regulation and oxidative response genes in the male and female genital tracts. (A) Expression levels of phage genes are shown on the x-axis and represent male urethral sample (blue) and female cervico-vaginal lavage (red). Expression levels of significantly differentially iron genes (B) and oxidative stress genes (C). * Indicates a gene more highly expressed in the male genital tract and # indicates a gene more highly expressed in the female genital tract (q-value of < 0.05). Red and blue X’s indicate the male and female counterparts of the dyad.

**Supplemental Figure 2.** Hamming distance tree and SNP distance matrix of *N. gonorrhoeae* isolates. Tree visualizing Hamming distances of total pairwise SNPs identified across strains. Scale bar shows distance of 300 SNPs.

**Table S1.** Deposited genomic data

**Table S2.** Characteristics of Female Subjects

**Table S3.** Top ten most highly expressed genes in each male subject

**Table S4.** Composite RPKM values for each gene in men *in vivo* vs *in vitro*

**Table S5.** Composite RPKM values for significantly different non-coding RNA genes in men *in vivo* vs *in vitro*

**Table S6.** Composite RPKM values for significantly different predictive sRNAs in men *in vivo* vs *in vitro* AND men *in vivo* vs women *in vivo*

**Table S7.** Composite RPKM values for each gene in men vs women

